# Tracking Transgenes with Color: RUBY as a Visual Marker in CRISPR-Edited Mutant Plants in Two Triticum Species

**DOI:** 10.64898/2026.02.10.705172

**Authors:** Rakesh Kumar, Ann Palayur, China Lunde, Ksenia V. Krasileva, Matthew J. Milner

## Abstract

CRISPR-Cas9 is a powerful tool for precise genome editing in plants, but the presence of foreign DNA, such as T-DNA, raises regulatory concerns and complicates mutant screening and field studies of edited material. Detecting plants with good transgene expression and later removing the T-DNA from edited plants is both time-consuming and costly. To address this, we developed a system that uses the non-destructive RUBY reporter, linked to the CRISPR-Cas9 cassette, and expressed under the *ZmUbi1* promoter. To assess the applicability of the system, it was tested on two *Triticum* species, targeting three genes in either tetraploid or hexaploid wheat. Strong correlations were observed in both T_0_ and T_1_ plants between betalain content and Cas9 expression, allowing for the quick identification of plants likely to be edited. Furthermore, the RUBY reporter could be used to select against the transgenic CRISPR-Cas9 cassette in subsequent generations at both the seed and seedling stages, thereby reducing the number of plants that need to be screened to identify edited lines without a T-DNA. This approach, using a nondestructive reporter, enabled rapid distinction between transgene expression in primary transgenics and served as a negative selection in the T_1_ generation, streamlining selection towards edited and T-DNA-free progeny.

## Introduction

Precise genome-editing methods, such as CRISPR/Cas, have revolutionized functional genomics and crop breeding by enabling efficient targeted genome modifications. The CRISPR/Cas system has gained popularity over other programmable nucleases (meganucleases, zinc-finger nucleases, and transcription activator-like effector (TALE) nucleases) due to its high specificity and ease of use. With CRISPR/Cas having a high level of adoption, it has also been readily adapted to perform higher-level genome modifications, including base editing, prime editing, and transcriptional enhancers/repressors, thereby broadening its applicability (Gilbertson *et al*., 2025). In a little more than a decade, CRISPR systems have undergone significant evolution and are now widely used in plants, particularly in crops, to target genes associated with traits of agronomic importance such as biotic and abiotic stress, nutrient uptake, male sterility, plant morphology as examples (Ibrahim *et al*., 2022; Kan *et al*., 2023; Li *et al*., 2022; Milner *et al*., 2020a).

Despite these advances, one persistent bottleneck in crop genome editing remains the need to quickly identify edited primary transformants and quickly remove the editing components. The ability to select plants with high transgene expression often involved measuring either transcript or protein levels to identify highly expressing plants. This is very important in polyploid species such as wheat where multiple copies of genes typically need to be knocked out to observe a phenotype.

Once the targets are edited, there is a desire to quickly remove the CRISPR/Cas components and obtain transgene-free edited plants without requiring multiple generations which could contribute to off-target effects (Aliaga-Franco *et al*., 2019; He *et al*., 2018). The traditional segregating or crossing approach is a time-consuming method for eliminating foreign DNA. In polyploid species such as wheat, this is further complicated by factors such as a higher number of complete or partial T-DNA inserts and genotyping costs, making it a lengthy and financially burdensome process to obtain edited material free of a transgene (He *et al*., 2018; Huang *et al*., 2023). In wheat, it takes approximately four months to generate transgenics with a CRISPR-Cas cassette and target-specific gRNA(s) (Smedley *et al*., 2021), and typically requires two to three generations to remove transgenes by Mendelian segregation from the edited plants. There is a pressing need to establish a rapid, high-throughput mutant screening system in the early stages of plant development to reduce the number of plants required to screen to obtain the desired mutants.

Traditionally, plant transformation relies on selectable markers such as antibiotics (e.g., hygromycin, kanamycin) or herbicide resistance genes to identify plants carrying a T-DNA (e.g., bar, cp4-epsps) (Chen *et al*., 1998; Fraley *et al*., 1983; Funke *et al*., 2006; Ortiz *et al*., 1996). Although these markers are effective, they have certain limitations: they lack non-destructive direct visual cues, may place metabolic burdens on the system, and could hinder downstream regulatory approval or consumer acceptance and do not give a quantitative measure of expression (Chen *et al*., 2018; Huang *et al*., 2023).Alternative reporter systems, such as GFP, GUS, and DsRED, are also sometimes used; however, they often require specialized equipment, destructive testing, or fail to produce strong signals across all tissues (He *et al*., 2020). A reliable, quantifiable, and readily observable marker that directly indicates the presence and expression of transgenes would significantly accelerate the detection of high-quality transgenic events in genome-editing workflows.

The RUBY reporter system provides an effective method for visually and non-destructively detecting betalain pigments. The *RUBY* reporter comprised a synthetic set of genes, *CYP76AD1, DODA*, and a glucosyltransferase (*GT*), that function as a polycistronic cassette, encoding a three-gene pathway to produce the visual reporter of a ruby color from betalain (He *et al*., 2020). The three betalain biosynthetic genes can be fused into a single open reading frame by adding 2A peptides, allowing them to undergo self-cleavage and release the individual enzymes for betalain biosynthesis between each ORF, which can be expressed under a single promoter and terminator (Ahier and Jarriault, 2014). These three enzymes convert endogenous tyrosine found in every cell into betalain pigments visible to the naked eye. Transgenic plants expressing *RUBY* produce a red-to-violet color that is clearly visible across different tissues without staining, microscopy, chemical assays, or tissue destruction. *RUBY* has been effectively used across various plant species, including wheat, as a simple visual marker for transformation and for monitoring transgene expression (Chen *et al*., 2024; He *et al*., 2020; Lee *et al*., 2023; Prusty *et al*., 2025).

In many polyploid species, strong expression of the *Cas9* gene is essential for editing multiple loci simultaneously in a single plant (Milner *et al*., 2020b; Wang *et al*., 2022). We hypothesize that RUBY can serve as a visual marker to distinguish transgenic plants that highly express *Cas9*, when further linked via a P2A peptide to the RUBY reporter. In the T_0_ generation, plants can be quickly selected to identify those highly expressing *Cas9*, and then selected against in subsequent generations to more quickly and economically recover non-transgenic genome-edited mutant plants. We evaluated the performance of this strategy in durum and hexaploid wheats, examined the correlation between RUBY pigmentation and transgene presence, and investigated cases where *RUBY* DNA sequences were present but pigment accumulation was absent. Our results demonstrate the potential of RUBY as a simple, cost-effective marker for determining transgene presence or absence in CRISPR-edited wheat plants, thereby streamlining crop genome-editing pipelines.

## Results

To understand if a visual reporter could serve as an indicator of transgene expression, the constitutive promoter maize ubiquitin (*ZmUbi1*) was used to drive expression of *Cas9* linked to the *RUBY* cassette via an additional P2A self-cleaving peptide. This configuration enabled the simultaneous expression of the four coding sequences under a single strong constitutive promoter. The *Cas9-RUBY* construct was introduced into two separate *Triticum* species via Agrobacterium-mediated transformation to test its efficacy as a non-destructive reporter and to serve as a direct visual marker for the presence and expression of *Cas9* (Figure 1A).

**Figure 1:**
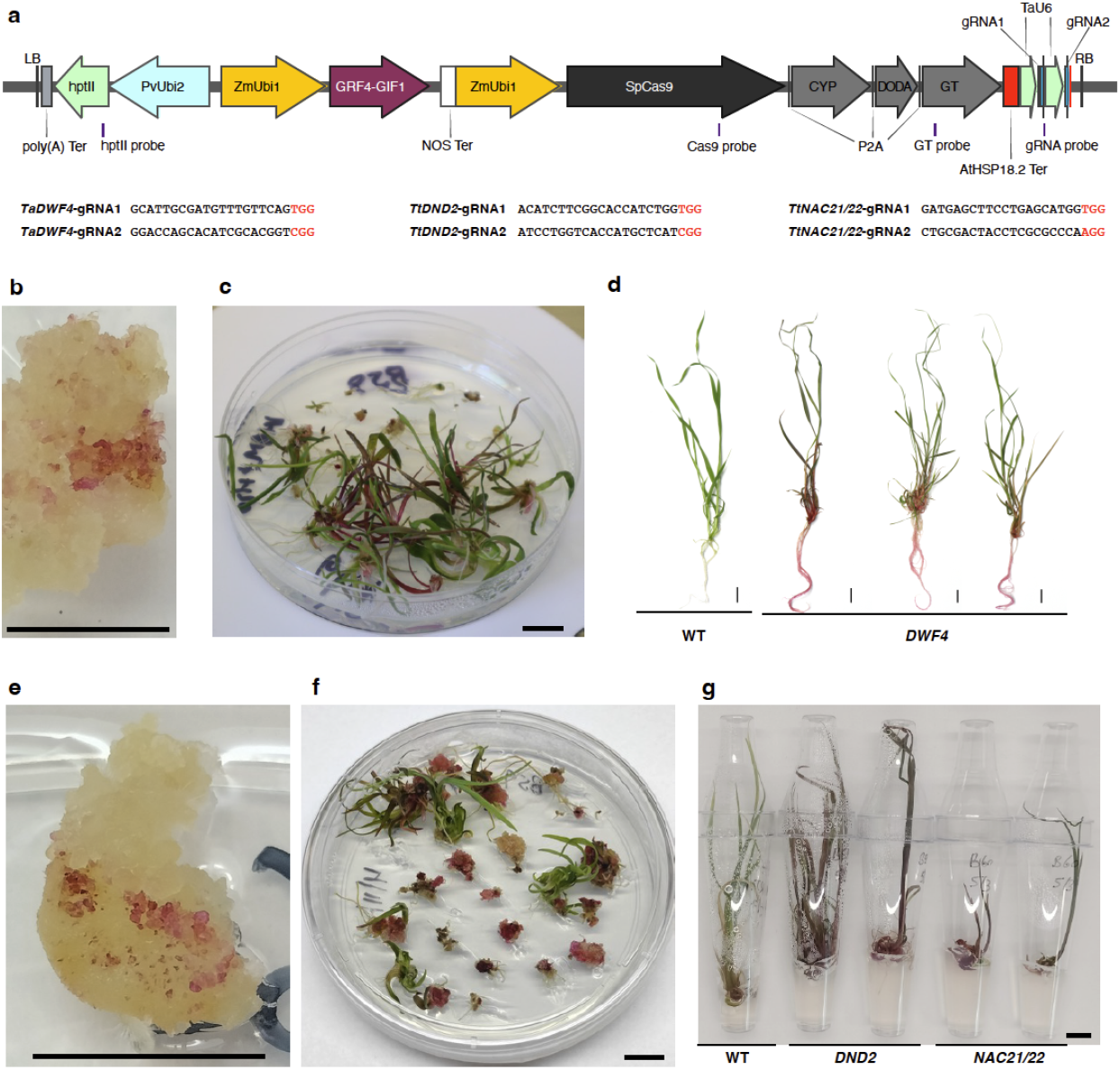
Design and validation of the Cas9-P2A-RUBY genome-editing and reporter system in Fielder and Kronos for editing four target genes (*TaDWF4* in Fielder, *TtDND2-1, TtDNA2-2*, and *TtNAC21/22* in Kronos). (a) Schematic of the T-DNA used for simultaneous genome editing and visual RUBY reporting. The construct contains *hptII* for selection, *TaGRF4-GIF1* to enhance regeneration, and a ZmUbi1-driven *SpCas9-P2A-CYP-DODA-GT* cassette that enables both CRISPR editing and betalain biosynthesis. Two gRNAs expressed under TaU6 promoters were used to target each gene. Red-highlighted nucleotides indicate the PAM site for the corresponding gRNA. Probe positions for *hptII, Cas9, GT*, and gRNA used in ddPCR copy-number assays are indicated. Guide RNA target sequences are shown below the construct map. (b-d; Fielder) and (e-g; Kronos): Embryogenic callus with red betalain sectors marking successful T-DNA delivery. Regenerated plantlets on selection medium exhibit strong RUBY pigmentation and display representative regenerated shoots and roots with characteristic red coloration, allowing rapid visual identification of transgenic events during tissue culture and regeneration. Scale bar for (b) and (e) = 0.4 cm; otherwise = 1 cm.

Ruby color was clearly visible at the callus induction and regeneration stages of tissue culture when driven by the *ZmUbi1* promoter (Figure 1B-G), confirming the expression of all four CDSs of this hybrid construct in both wheat backgrounds during plant transformation. No tissue culture abnormalities were seen with the *Cas9-RUBY* construct in either wheat species. At later developmental stages, regenerated plantlets exhibiting a range of ruby coloration were advanced for further analysis.

To determine whether the coloration of transgenic Fielder plants matched *Cas9* expression, expression of all four genes was measured and compared with betalain content in 24 plantlets. qPCR-based expression analysis revealed that some green plants expressed all four CDSs of the hybrid construct but did not develop detectable betalain in the leaves (Figure 2). The correlation (Spearman’s rho) between *Cas9* expression and the CDS units of the *RUBY* reporter genes (*CYP76AD1*, 0.89; *DODA*, 0.88; and *GT*, 0.90; p < 0.001) was strong, confirming functional validation of expression of four CDSs under the same promoter. Expression of all four CDSs showed strong positive correlations with betalain production (*Cas9*, 0.78; *CYP76AD1*, 0.74; *DODA*, 0.82; and *GT*, 0.78; all p<0.001). We also observed that some transgenic plantlets exhibited red color only in their shoots, others only in their roots, and others throughout the entire plantlet (Figure S1).

**Figure 2:**
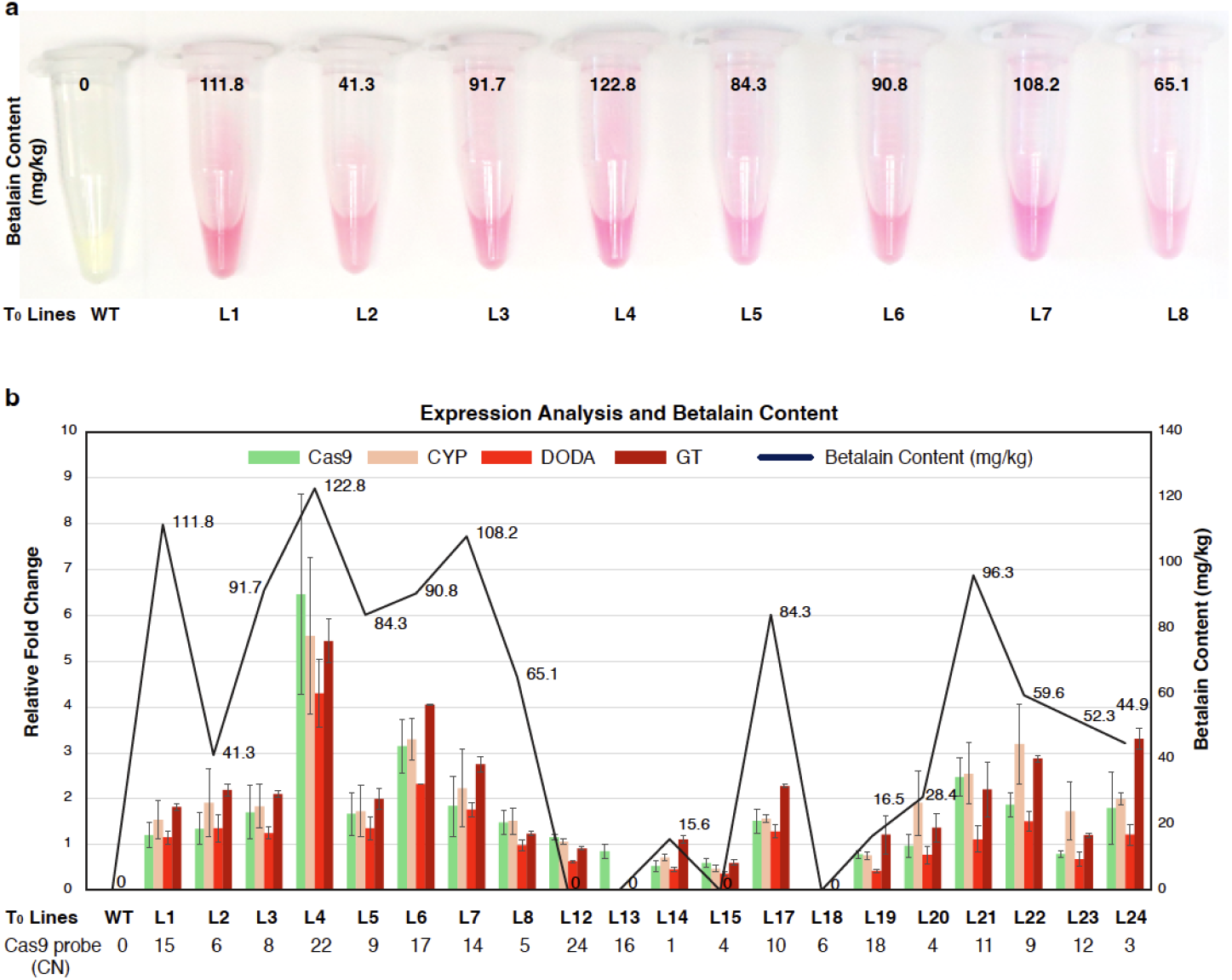
Transgene dosage–dependent expression of the RUBY genes and betalain accumulation in independent Fielder T_0_ Cas9-RUBY events. (a) Visual comparison of betalain extracts from wild type (W) and eight independent lines (1–8). Tubes show variation in pink/red pigmentation corresponding to different levels of betalain production. (b) Integrated expression, copy-number, and metabolite profiling of a larger set of independent lines. Bar plots show relative expression of *Cas9* (green), *CYP* (peach), *DODA* (red), and *GT* (dark red), while the blue line indicates betalain content (mg/kg fresh weight). *Cas9* copy number (CN), measured by ddPCR, is listed below the x-axis.

To understand which factors could influence the levels of betalain, correlations (Spearman’s rho) between the T-DNA copy number and expression were measured. The copy number of the different genes of the *Cas9-RUBY* construct was measured in both Fielder and Kronos T_0_ plantlets (Figure 1A). From the ddPCR data, it was observed that the T-DNA was inserted in a fragmented manner, with varying copy numbers from left border (LB) to right border (RB) (Figure 3, Table S3-5). Overall, relative to the *hptII* probe designed from the *hptII* located near to the LB in the T-DNA, single-copy insertions were not observed in Fielder, whereas 12% and 31% single-copy insertions were observed in Kronos for targeted knockouts of *TtDND2-1* and *TtDND2-2*, or of *TtNAC21/22*, respectively. In both species, we observed more insertions/higher copy numbers with probes located near the RB than with probes located near the LB, indicating incomplete and concatemerized insert transfer.

**Figure 3.**
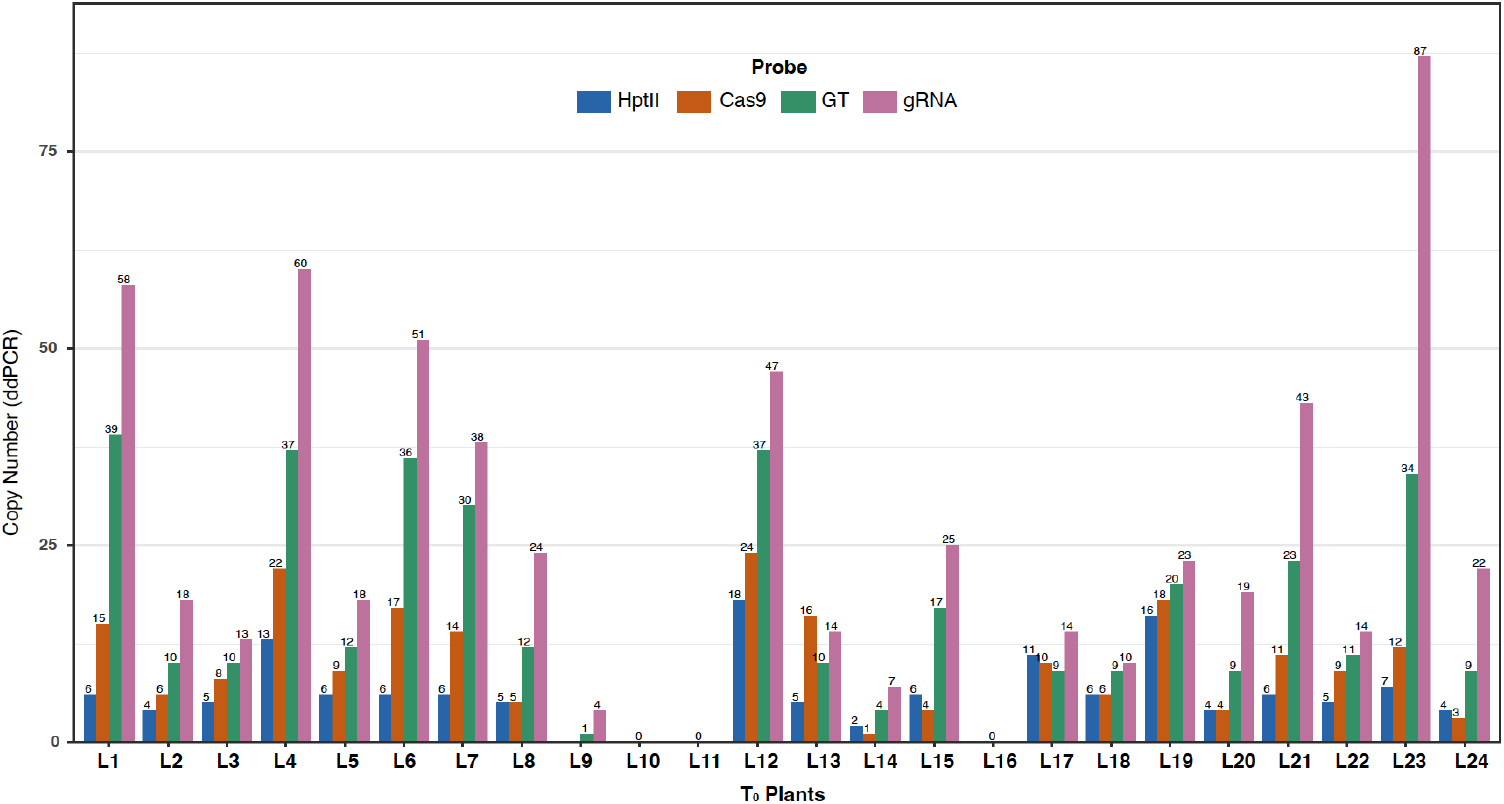
Distribution of T-DNA inserts assessed by ddPCR in independent T_0_ Fielder wheat lines. Bar plots show copy numbers for four T-DNA components (*hptII, Cas9, GT*, and gRNA) measured by ddPCR across 24 independent T_0_ Fielder events (lines 1–24). Each bar corresponds to one of the four probes, and the values are indicated above it.

**Figure 4:**
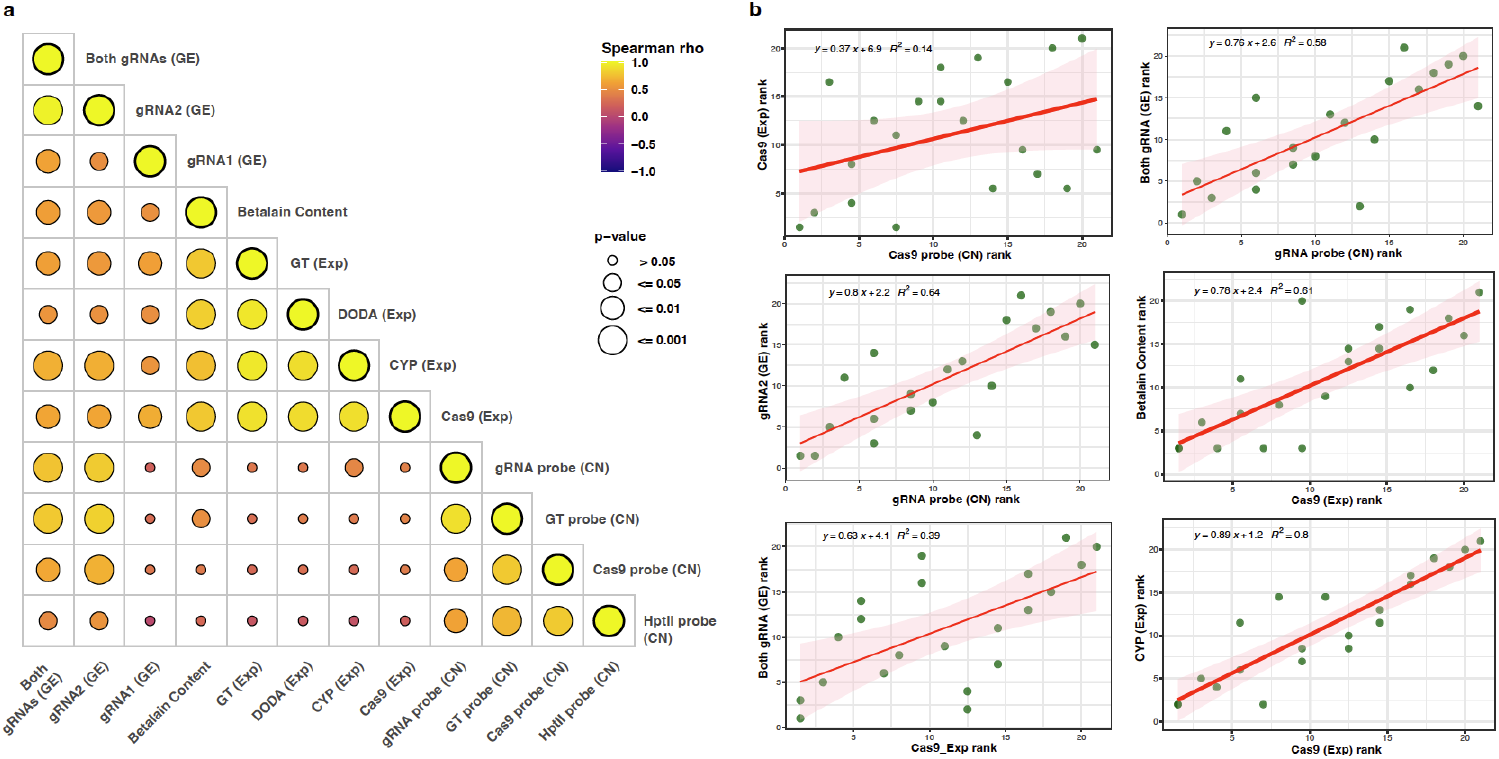
Integrated relationships among copy number, expression, and genome editing underlying RUBY betalain accumulation in Fielder. (A) Spearman correlation matrix summarizing pairwise associations among T-DNA copy number (CN) for *hptII, Cas9, GT*, and gRNA probes; transgene expression (*Cas9* (Exp), *CYP* (Exp), *DODA* (Exp), and *GT* (Exp)); genome-editing outputs (gRNA1 (GE), gRNA2 (GE), and both gRNAs (GE)); and betalain content. Circle color represents the Spearman correlation coefficient (ρ; −1 to +1), while circle size indicates statistical significance (p-value thresholds shown at right). (B) Rank-based regression analyses validating key relationships from panel A.

In Fielder, there was a strong correlation among the copy numbers of the various regions of the T-DNA with strong correlations observed from probes located near or next genes on the T-DNA, (*hptII* and *Cas9* probe, 0.79; *Cas9* and *GT* probe, 0.79; *GT* and gRNA probe, 0.90; p <0.001), the correlation between the probe closest to the LB (hptII probe) and RB (gRNA probe) was the lowest at 0.61 but a significant correlation between the copy numbers was still observed (p<0.01) (Table S6). It was also observed that the copy number of the *Cas9* region was not significantly correlated with *Cas9* expression (0.37), *CYP76AD1* expression (0.25), *DODA* expression (0.29), or *GT* expression (0.25), all with p-values > 0.05, suggesting substantial positional effects. To assess the correlation between *Cas9* expression and the *RUBY* reporter and overall editing rates, mutations were identified in the T_0_ lines. Of the 24 putative T_0_ *Cas9-RUBY* plants targeting *TaDWF4*, 20 plants were confirmed to have been edited at least one of the two gRNA regions for at least one of the three *TaDWF4* homeologues. As expected, the plants not containing a T-DNA (Lines 10, 11, and 16) and the partial T-DNA-containing event (Line 9) did not show any editing at either gRNA. Expression of *Cas9* was a strong predictor of editing, with strong correlations between expression of any of the four genes and editing at either gRNA site or at both sites combined (p< 0.01). Interestingly, a strong correlation was also observed between gRNA probe copy number and editing at gRNA2 (0.80; p<0.001) of the *TaDWF4* gene, suggesting that higher gRNA expression levels can also improve editing. On the other hand, the correlation between gRNA probe copy number and editing at gRNA1 was weak and not statistically significant.

To determine whether the similar correlations observed in Fielder held in tetraploid wheat, a set of Kronos T_1_ lines targeting *TtDND2-1 and TtDND2-2* or *TtNAC21/22* were selected based on their copy number measured at the *Cas9* region of the T-DNA. Expression analysis and betalain content measurements of the Kronos T_1_ plants revealed that the copy number of the *Cas9* probe again showed no correlation with the overall expression of the *Cas9* transgene (Figure 5, Figure S2, Table S7). However, the expression of four CDSs in the hybrid construct was again significantly correlated with betalain content (*Cas9*, 0.83; *CYP76AD1*, 0.83; *DODA*, 0.74; and *GT*, 0.74; p < 0.05). Furthermore, the expression of three CDS genes of the *RUBY* reporter showed a very strong correlation (*CYP76AD1*, 1.0; *DODA*, 0.98; and *GT*, 0.98) with *Cas9* expression (p<0.001), which again confirmed the functionality of the four CDS units under the same promoter in tetraploid wheat. The correlations between *hptII* and *Cas9* probes, *Cas9* and *GT* probes, and *GT* and *hptII* probes were 0.85 (p-value <0.01), 0.94 (p-value <0.001), and 0.74 (p-value <0.05), respectively. Additionally, the T-DNA copy number in these seedlings was determined in parallel with the editing status (Table S8-10). The data suggested that, overall, both *TtNAC21/22* gRNAs were more efficient at creating edits than the *TtDND2* gRNAs; however, no correlation between editing and *GT* probe copy number was observed in Kronos (Table S8).

**Figure 5:**
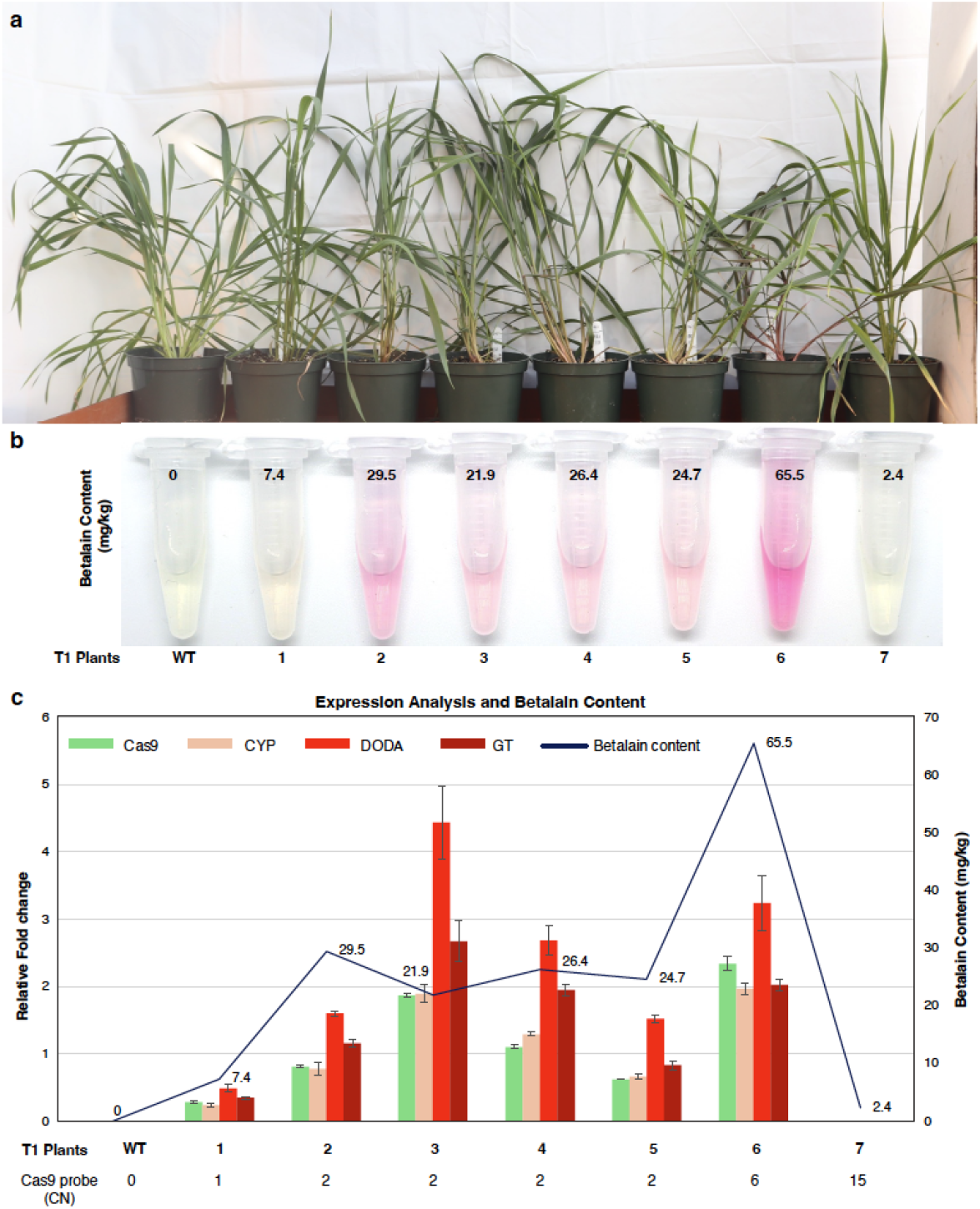
Variation in RUBY transgene dosage, expression, and betalain pigmentation among Kronos T_1_ plants. (a) Phenotypic comparison of representative T_1_ plants (wild type control (WT) and lines 1–7 (NAC21/22_L7-1, NAC21/22_L4-1, NAC21/22_L6-5, DND2_L6-2, DND2_L7-1, DND2_L2-5, DND2_L5-7) carrying the *Cas9-RUBY* reporter, shown as whole plants and (b) corresponding betalain extracts. Numbers above each tube indicate quantified betalain content (mg/kg fresh weight) in T_1_ events. (c) Integrated analysis of transgene expression and betalain accumulation in the same Kronos T_1_ plants. Bar plots show relative expression levels of *Cas9* (green), *CYP* (peach), *DODA* (red), and *GT* (dark red), while the blue line denotes betalain content (mg/ kg). *Cas9* copy number (CN), determined by ddPCR and listed below the x-axis.

To determine whether one could select against RUBY-colored plants in the next generation, transgenic Fielder T_1_ seeds were harvested and sorted by eye based on observed ruby color in the grains (Figure 6). Coloration of the grain was not uniform in seeds harvested from Fielder plants expressing RUBY (Figure S2). The strongest ruby coloration was on the ventral side of the grain around the crease and on the dorsal side containing the embryo. Whereas coloration on the dorsal side of the grain distal to the embryo, was generally weaker (Figure S2). To understand if grain coloration could be used to select for lower copy number plants in the next generation, eight seeds showing the most ruby coloration and the eight seeds showing the least ruby coloration were germinated from five independent T_0_ lines. Initial observations showed no obvious ruby color in the developing seedlings; however, three weeks after planting, ruby color was observed in the coleoptile (Figures 6, Figure S3). We observed that the ruby color became visible in other tissues over time, beginning in the stem and progressing apically. Emergent leaves were always green.

**Figure 6:**
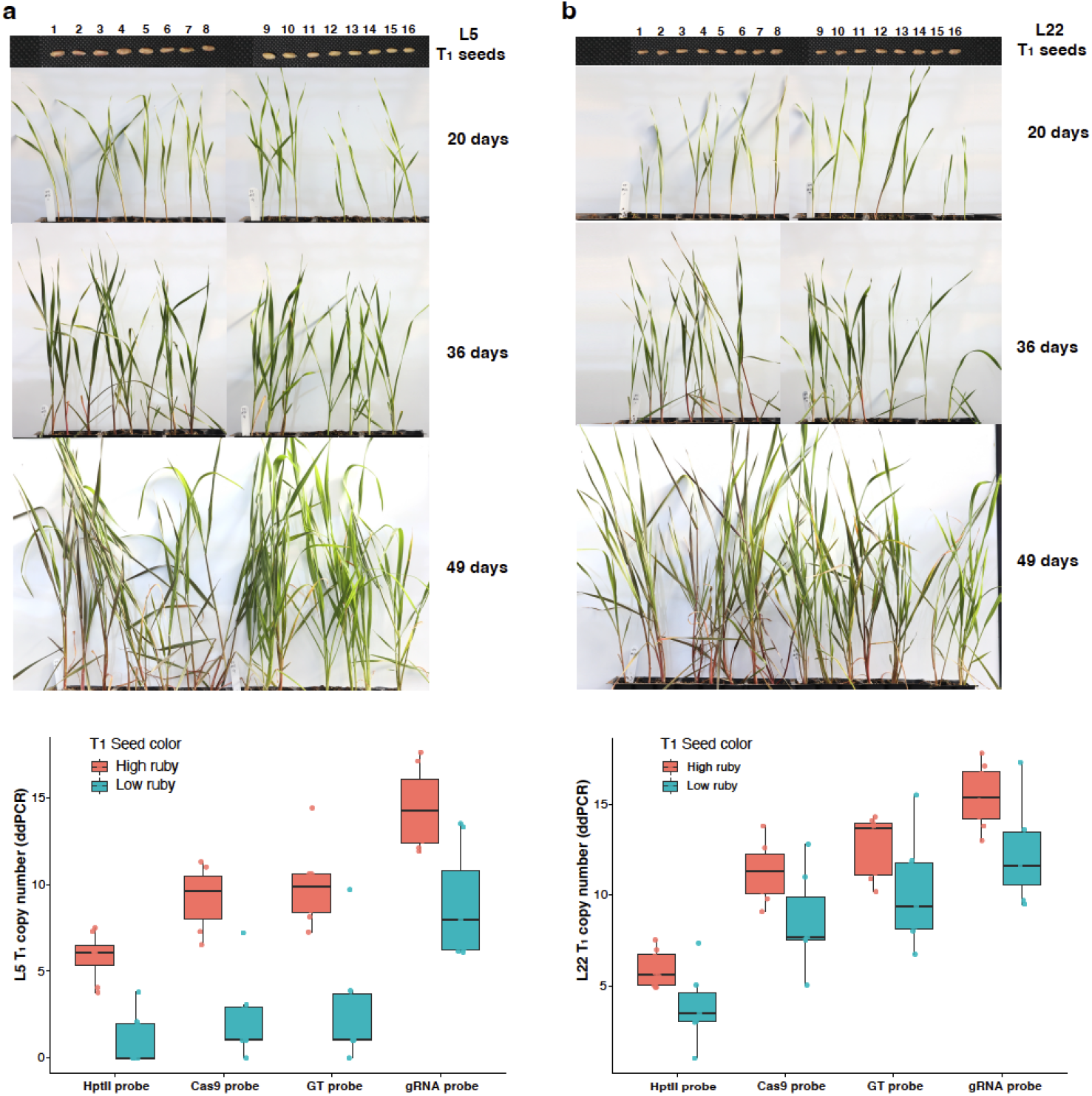
RUBY-dependent segregation and copy-number variation in two representative T_1_ Fielder wheat lines. (a-b) Two independent T_1_ families (L5 in panel a and L22 in panel b) are shown, each derived from a single T_0_ Fielder line. For each line, 16 T_1_ seeds are displayed at the top, grouped by seed-coat ruby intensity into high-ruby (1-8) and low-ruby (9-16) categories. Corresponding seedlings were grown and imaged at 20, 36, and 49 days after sowing, showing betalain pigmentation accumulation in older tissues, particularly in the stems and base of the leaf. The boxplots at the bottom summarize ddPCR-based T_1_ copy-number distributions for *hptII, Cas9, GT*, and gRNA probes, comparing plants grown from high-ruby (red) and low-ruby (teal) seed pools for each line.

To understand how well one could select against potential T-DNAs in the T_1_ generation using seed color as a marker, the copy number of the T-DNA and edits were measured in germinated seedlings (Table S11). In all five T_1_ lines tested, plants derived from high colored ruby seeds contained, on average, higher copy numbers of all T-DNA components, whereas plants derived from low-ruby colored seeds contained, on average, lower-copy insertions for each region of the T-DNA (Figure 6). In one of the five lines tested, plants were recovered without any full-length T-DNAs in the T_1_ generation. Although small fragments of the RB (gRNA probe) were still present. This demonstrates that seed-coat RUBY intensity reflects transgene dosage in the T_1_ generation and can be used to select against it. There were also instances of high-copy-number plants lacking ruby coloration, suggesting large positional effects of the inserted T-DNAs and multiple insertion sites in the lines tested.

Sequencing analysis of T_1_ plants indicated the presence of new mutations not observed in T_0_ indicate functional activity of the ribonucleoprotein (RNP) complex at later developmental stages of T_0_ or during the initial phase of T_1_ seedling development (Table S11). In the *TaDWF4* T_1_ lines, L2, L3, L5, L17, and L22 showed editing rates of 100%, 100%, 85.7%, 87.1%, and 100%, respectively, with either gRNA, across at least one DWF4 homeologue gene. Comparison to T_0_ plants, showed maximum editing of 42.1%, 40%, 33.3%, 8.3% and 84.7% at any given homeologue for *TaDWF4* in L2, L3, L5, L17 and L22, respectively (Table S3). It should also be noted that out of the high or low ruby color seed categories in the five lines tested, a given offspring was not more or less likely to be edited in the T_1_ generation based on the level of color observed in the grain.

## Discussion

CRISPR-based genome editing is a promising approach to achieving more resilient crops. Diverse strategies have been employed to generate genome-edited plants, including transient expression using an RNP complex, additive shoot inoculation, grafting, virus-mediated transformation, and *Agrobacterium*-mediated transformation via stable integration of the CRISPR components into the plant genome (Liang *et al*., 2017; Maher *et al*., 2020; Yang *et al*., 2023). While these approaches have enabled substantial progress, each has limitations, including species-specificity, inefficient delivery, and the transgenerational persistence of viral vectors. Among these methods, *Agrobacterium*-mediated transformation remains the most efficient and widely available approach for many agricultural crops, particularly cereals such as wheat. However, stable integration of foreign DNA into the plant genome raises concerns related to biosafety, regulatory acceptance, and gene flow (Chen *et al*., 2018; Huang *et al*., 2023; Tuteja *et al*., 2012). These concerns underscore the need for tools that facilitate the rapid identification of edited, transgene-free plants in subsequent generations. RUBY has emerged as a versatile visual marker that enables non-destructive identification of transgenic tissues without specialized equipment and chemical assays (He *et al*., 2020). It has been successfully employed across diverse plant species for various purposes, ranging from transformation optimization to environmental stress assessment (Adhikari *et al*., 2025; Jeong *et al*., 2024; Prusty *et al*., 2025; Yu *et al*., 2023). To understand if *RUBY* could be used as both a positive and negative selection marker, we analyzed the integration of the synthetic betalain biosynthesis pathway. *RUBY* and *Cas9* were linked and expressed under a common promoter, to detect high-quality events and assess the likelihood of editing in these plants. The *Cas9-RUBY* fusion construct driven by the maize ubiquitin promoter (*ZmUbi1*) ensured strong coordinated expression of *Cas9* and the *RUBY* marker. Previous reports have shown that high *Cas9* expression is paramount for achieving high editing levels in wheat (Milner *et al*., 2020b). In our *Cas9-RUBY* construct, the P2A peptide enables production of functional Cas9 and RUBY proteins from a single transcript. Thus, betalain content was strongly associated with *Cas9* expression and ensured the presence of transcriptionally active transgenes. In the two *Triticum* species, the plantlets expressing *Cas9*, and all three *RUBY* CDSs confirmed the functionality of the hybrid construct and showed that other plant species may also benefit from similar approaches. Molecular analysis of the target genes confirmed editing in multiple loci, indicating that the RUBY P2A fusion does not impair Cas9 activity and editing efficiency. However, we found that as many other have noted that editing efficiency at some loci (*TaDWF4* locus using gRNA1 in Fielder) was lower, even in plants strongly expressing *Cas9* and *RUBY*, indicating that factors beyond *Cas9* expression such as gRNA sequence, promoter selection, and local chromatin environment, may also affect overall editing efficiency (Kor *et al*., 2022; Liang *et al*., 2016; Liu *et al*., 2019; Milner *et al*., 2024).

It was also observed that T_0_ transgenic wheat plants generated by *Agrobacterium*-mediated transformation frequently harbor multiple T-DNA inserts, with high copy numbers of partial T-DNA sequences rather than simple single-copy complete insertions. This was true in both tetraploid and hexaploid wheats. The observed pattern and segregation of the T_1_ plants strongly suggest that T-DNA molecules undergo concatemerization prior to chromosomal integration, rather than independent insertion events at multiple loci, as large steps in copy number were observed rather than in discrete units. Similar complex insertion architectures have been reported in wheat (Wu et al., 2006) and maize (Neelakandan et al., 2023) and are consistent with models of T-DNA integration that involve host DNA repair pathways, such as non-homologous end joining (NHEJ) and microhomology-mediated end joining (MMEJ). This complicates the ability to quickly clear the T-DNAs when multiple T-DNA fragments lacking a reporter are present in the transformant. Thus, multiple markers may be required to select for strong expression and clear RB fragments in subsequent generations. Or at least place these markers at the RB, as this is the most frequent insertion in the *Triticum* species tested.

While others have used similar methodologies to quickly select plants that remove T-DNAs in other plant species (Aliaga-Franco *et al*., 2019; He *et al*., 2018). These previous reports help clear the plants of the transgene, such as the RUBY system, but do not identify plants showing strong Cas9expression. These systems are unable to quickly identify high-expressing plants because the systems are seed-based. Future systems that combine positive selection (e.g., RUBY) with negative selection (e.g., barnase) may further improve the generation of novel mutants that rapidly eliminate editing components in subsequent generations. As a typical T_0_ plant with multiple T-DNA loci may contain inserts in both euchromatic and heterochromatic regions with at least one copy, typically including the selectable marker, must be transcriptionally active for recovery under selection in tissue culture, other copies may be transcriptionally silent due to positional effects or epigenetic silencing, further complicating the removal of unwanted DNA in subsequent generations (Thomson *et al*., 2024). We observed transgenics in the advanced T_1_ lines, with some plants having higher copy number but lower bet alin production than the parental plant (Figure 6). Consequently, some T_1_ progeny may lack the phenotypic effects of the transgenes even when partial or fragmented T-DNA remains in the genome. This underscores the continued need for comprehensive molecular screening across the entire T-DNA (LB to RB) before labeling a plant as transgene-free.

A range of various RUBY expression patterns in T_0_ and T_1_ plants were observed, including pigmentation limited to shoots, roots, or the entire seedlings. Our findings emphasize that even robust constitutive promoters such as ZmUbi1 do not ensure consistent or persistent transgene expression across tissues or developmental stages. Reliance on visual markers alone may yield false negatives, particularly when tissue-specific promoters are used or when transgene expression is influenced by chromatin context or epigenetic modifications.

However, using the *Cas9-RUBY* fusion system in gene editing transgenic plants can reduce the number of plants required to screen to obtain the desired edits by quickly identifying plants most likely to be edited. This same marker can then be used to reduce the number of subsequent generations needed to obtain transgene-free edited lines, thereby lowering costs, labor, and time in plant genome-editing pipelines.

## Materials and Methods

### Binary Vector Constructions and gRNA Design for Target Genes

The binary vectors and DNA fragments used in this study are listed in Table S1. For wheat transformations, the vector JD633 (Addgene #160393) was modified to generate pLRK16 by linking the *RUBY* gene with *Cas9* using P2A (wheat codon optimized) immediately after the C-terminal NLS signal. To construct pLRK16 a fragment of the *Cas9* gene containing the N-terminal NLS signal was synthesized (Integrated DNA Technologies, USA) along with the additional three fragments containing the C-terminal part of the Cas9 gene with a P2A peptide, the complete *RUBY* gene, *AtHSP18*.*2* terminator, and a portion of the *TaU6* promoter (Table S1). The amplified and synthesized fragments were ligated into the SbfI and PmlI-digested JD633 vector using the NEBuilder® HiFi DNA Assembly Master Mix (New England Biolabs) to create pLRK16. To allow for two gRNAs targeting each gene, a further fragment containing gRNA1_RNAScaffold_PolII terminator-TaU6 promoter_gRNA2 was synthesized (Integrated DNA Technologies, USA) to accommodate two gRNAs for each target gene and driven by independent TaU6 promoters (Table S1). The synthesized fragments were flanked with PaqCI sites for guide sequences to be added.

To perform targeted knockout of *TaDWF4* (TraesCS4A02G078000, TraesCS4B02G234100, TraesCS4D02G235200), *TtNAC21/22* (TrturKRN5A02G008060, TrturKRN5B02G010390), and *TtDND2-1* (TrturKRN1A02G046790, TrturKRN1B02G056340), *TtDND2-2* (TrturKRN3A02G044450, TrturKRN3B02G052750)) were selected for targeting the wheat genomes. Potential gRNAs were identified using WheatCRISPR (https://crispr.bioinfo.nrc.ca/; Cram *et al*., 2019) to target all homeologues. Further, the gRNAs were checked for any off-target match in the genome using the BLAST tool available at EnsemblPlants (https://plants.ensembl.org/Triticum_aestivum/Info/Index) or the offline NCBI BLAST+ (Camacho *et al*., 2009) using the Kronos genome (Seong *et al*., 2026).

The final fully sequenced vectors were transformed into the chemically competent *Agrobacterium* strain AGL-1 using the heat-shock method (Höfgen and Willmitzer, 1988). The positive colonies were confirmed by colony PCR using binary plasmid backbone-specific primers (Table S2) before plant transformation. The sequence files of the verified binary plasmids are provided as supplementary files.

### Wheat Transformation

Hexaploid (*Triticum aestivum* L. cv ‘Fielder’ (CItr 17268)) and tetraploid (*Triticum turgidum* subsp. *durum* cv. ‘Kronos’ (PI 576168)) wheat genotypes were used for the *Agrobacterium*-mediated transformation of the plasmids pLRK57 (Fielder, *TaDWF4*), pLRK35 (Kronos, *TtDND2*), and pLRK36 (Kronos, *TtNAC21/22*). The method used to transform both wheat species was previously described by Hayta *et al*. (2019). Briefly, wheat plants were grown in a glasshouse until spikes had matured to 16-22 days after anthesis. The immature embryos from spikes were isolated from sterilized seeds under aseptic conditions in a laminar airflow. Immature embryos were inoculated with the *Agrobacterium* culture containing one of the plasmids above at an OD_600_ of 0.8 for three days in the dark at 21 ± 1 °C. The embryogenic axes were removed after three days and subcultured for subsequent tissue culture steps to obtain regenerated plantlets. The plantlets were hardened in soil in the seedling tray by covering the tray with a plastic dome for 5 days at 21 ± 1 °C under light with a 16/8-hour photoperiod. After acclimatization, individual plantlets were transferred to 6-inch plastic pots and grown in the glasshouse until maturity at 24-26 °C with a 16/8-hour photoperiod.

### Molecular Analysis and Copy Number Determination

Genomic DNA was extracted from the putative T_0_ transgenics using a modified CTAB method (Doyle, JJ and Doyle, JL, 1987). A microcentrifuge tube containing approximately 100 mg of leaf sample, along with two steel beads (0.45 mm), was snap-frozen in liquid nitrogen and immediately ground to a fine powder using a tissue grinder. The sample was mixed well after adding 600 μL of CTAB DNA extraction buffer (2% CTAB, 20 mM EDTA, 100 mM Tris-HCl, pH 8.0, 1.4 M NaCl, 0.5% sodium bisulfite (Na_2_O_5_S_2_), 0.1% β-mercaptoethanol) and incubated for 30 min at 65 °C in a water bath with intermittent mixing. The samples were cooled at room temperature for 10 min, then 600 μL of chloroform: isoamyl alcohol (24:1) was added, mixed briefly, and centrifuged at 10,000 × g for 15 min. The supernatant (∼500 μL) was transferred to a new 1.5 mL centrifuge tube containing 2 μL of RNAse (10 mg/mL) and incubated at 37 °C for an additional 30 min. An equal volume (500 μL) of chloroform: isoamyl alcohol (24:1) solution was again added, mixed briefly, and then centrifuged at 10,000 × g for 15 min to further purify the sample. The supernatant (∼350 μL) was transferred to a new 1.5 ml microcentrifuge tube, and mixed with an equal volume of isopropanol, incubated at -20 °C for 30 min, and centrifuged at 10000 x g for 15 min at 4 °C. The DNA pellet was washed once with 70% ethanol after discarding the supernatant, then dried at 37 °C to remove residual ethanol. The pellet was redissolved in 0.1X TE buffer and quantified using a Nanodrop ND-1000 (Thermo Scientific) or DeNovix DS-11 Fx+ (DeNovix Inc.) to determine DNA concentration. gDNA was used to detect T-DNA transgenes and to assess editing at target gene loci using the primer pairs listed in Table S2. Editing at target gene loci was analyzed using Sanger sequencing and/or Oxford Nanopore sequencing (Native Barcoding Kit 24 V14 (SQK-NBD114.24) and PromethION R10.4.1 flow cells, Oxford Nanopore Technologies).

The copy number of the T-DNA was assessed using digital droplet PCR (ddPCR) (Collier *et al*., 2017). For ddPCR: 20 μL reactions were set up to digest 670 ng of gDNA with PspXI, and incubated overnight at 37 °C. A 25 μL ddPCR cocktail containing digested gDNA (100 ng), 250 nM of each primer pair for both the endogenous gene and transgene, 125 nM for each probe, ddPCR Supermix for Probes, no dUTP (Bio-Rad Laboratories) was prepared for each sample. Twenty μL of the PCR mix and 70 μL of Bio-Rad Droplet Generation Oil were loaded into the microcapillary droplet generator cartridge (Bio-Rad) at their specified locations, resulting in a final volume of 40 μL droplets, according to the manufacturer’s instructions. The droplets were transferred to a skirted Eppendorf PCR plate and sealed with pierceable foil (Bio-Rad). The ddPCR was run with the following conditions: denaturation at 95 °C for 10 min, followed by 40 cycles at 94 °C for 30 sec and 60 °C for 1 min. With a final incubation at 98 °C for 10 min. The PCR plate was transferred to a QX200 droplet reader (Bio-Rad), and the transgene copy number was measured using the QuantaSoft™ software (v1.6.6.0320; Bio-Rad) with default settings. The primers specific to endogenous genes and transgenes used are listed in Table S2.

### Gene expression analysis

qPCR was performed to determine the expression levels of four CDSs (*Cas9* and three genes comprising the RUBY reporter) driven by the *ZmUbi1* promoter. A well-developed 4th or 5th leaf tissue was selected for qPCR and betalain quantification. The leaf was divided vertically along the midrib into two halves: one for qPCR and the other for betalain quantification. The total RNA was extracted from the leaf samples using the RNeasy Plant Mini Kit (QIAGEN) according to the manufacturer’s instructions. The total RNA samples were treated with DNase I (QIAGEN) to remove DNA contamination before cDNA synthesis. The cDNA was synthesized from 500 ng of total RNA per sample from the leaf using the Protoscript II first-strand cDNA synthesis kit (New England Biolabs). PowerUp™ SYBR™ Green Master Mix (Applied Biosystems) was used to quantify RNA levels, with the wheat actin gene (Fielder, TraesFLD1B01G310300; Kronos, TrturKRN1B02G048790) (Kumar *et al*., 2020) serving as internal controls and Cas9_RUBY cassette qPCR primers (Table S2). The transcript levels were analyzed using the ^ΔΔ^Ct method.

### betalain Content Quantification

betalain content was determined by methanol extraction (Pramanik *et al*., 2024). The leaf samples were weighed and ground to a fine powder in liquid nitrogen. The extraction solution (a 1:1 mixture of water and methanol) was added to the powdered samples at a volume equivalent to 10 times the fresh weight of the sample. The samples were thoroughly mixed, collected into a 1.7 mL microcentrifuge tube, and centrifuged at 21000 x g for 5 minutes at room temperature. The supernatant was transferred to a new1.7 mL microcentrifuge tube and, if required, further diluted in ddH2O to achieve the desired volume for measurements. The final solution was measured at 538 nm using a spectrophotometer. The reaction was used to calculate the betalain content using the following formula (mg/kg):

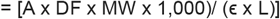

where:

- A, the absorption value at 538 nm.
- DF*, the dilution factor, 10 × sample dilution ratio.
- MW, the molecular weight of betalain, 550 g/mol.
- ϵ, the molar extinction coefficient, 60,000 L/mol.cm in H2O.
- L, the path length, 1 cm for 1 mL cuvette

### Statistical Analysis

Pairwise Spearman’s correlations were calculated using R for ddPCR copy number, betalain content, expression, and editing data in both wheat species (Dietrich and Leoncio, 2025). Rank-transformed data-based linear regressions were computed to measure the parameter relationships. The R-packages (“Hmisc” (Harrell, 2026), “ggplot2” (Wickham, 2016), “ggpubr” (Kassambara, 2025)) were used in RStudio to compute and draw the correlation plots.

## Author contributions

RK, KVK, and MJM conceived the idea. RK, AP, and CL performed experiments, collected data, and analyzed data. RK drafted the manuscript. RK, AP, CL, KVK, and MJM edited the manuscript.

## Funding

This project was funded by the USDA-NIFA award 2021-67013-35726, awarded to KVK as well as as part of the USDA project 2030-21430-015-000D awarded to MJM.

## Acknowledgements

We are grateful to Dr. Jennifer Bragg, Ms. Toni Mohr, Dr. Swarupa Nanda Mandal, and Dr. Shivani for their assistance with the analysis of the transgenic plants.

## Conflicts of Interest

The authors declare no conflicts of interest.

## Data Availability Statement

The data that supports the findings of this study are available in the Supporting Information of this article.

## Notes

### Competing Interest Statement

The authors have declared no competing interest.

