## Supplementary material for "Tracking Transgenes with Color: RUBY as a Visual Marker in CRISPR-Edited Mutant Plants in Two Triticum Species": Suppl Figures

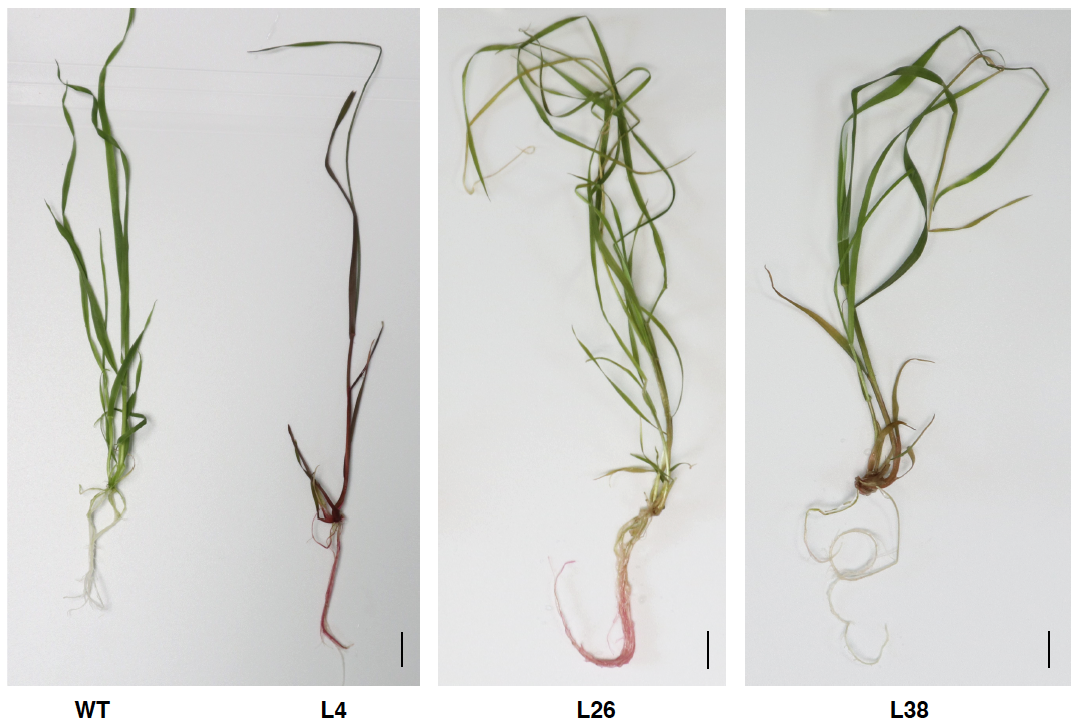

**Figure S1:** **Variations in RUBY color visibility in different parts of T_0_ plantlets in Fielder derived from Cas9-P2A-RUBY/gRNAs(DWF4) construct.** Scale bar = 1 cm.

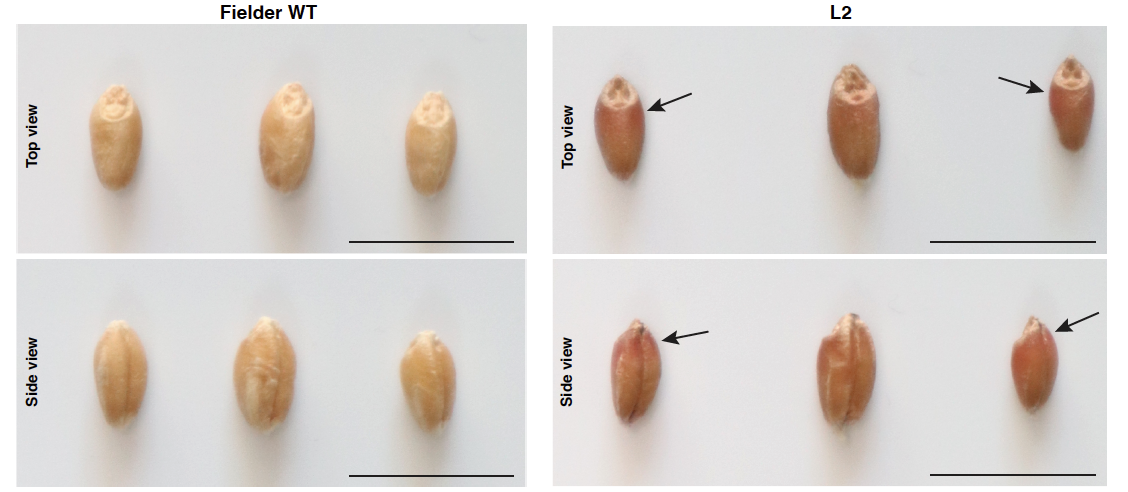

**Figure S2: A representation of localized variations in betalain accumulation in T_1_ seeds.** Fielder T_1_ seeds of L2 derived from the Cas9-P2A-RUBY/gRNA(DWF4) construct. Scale bar = 1 cm.

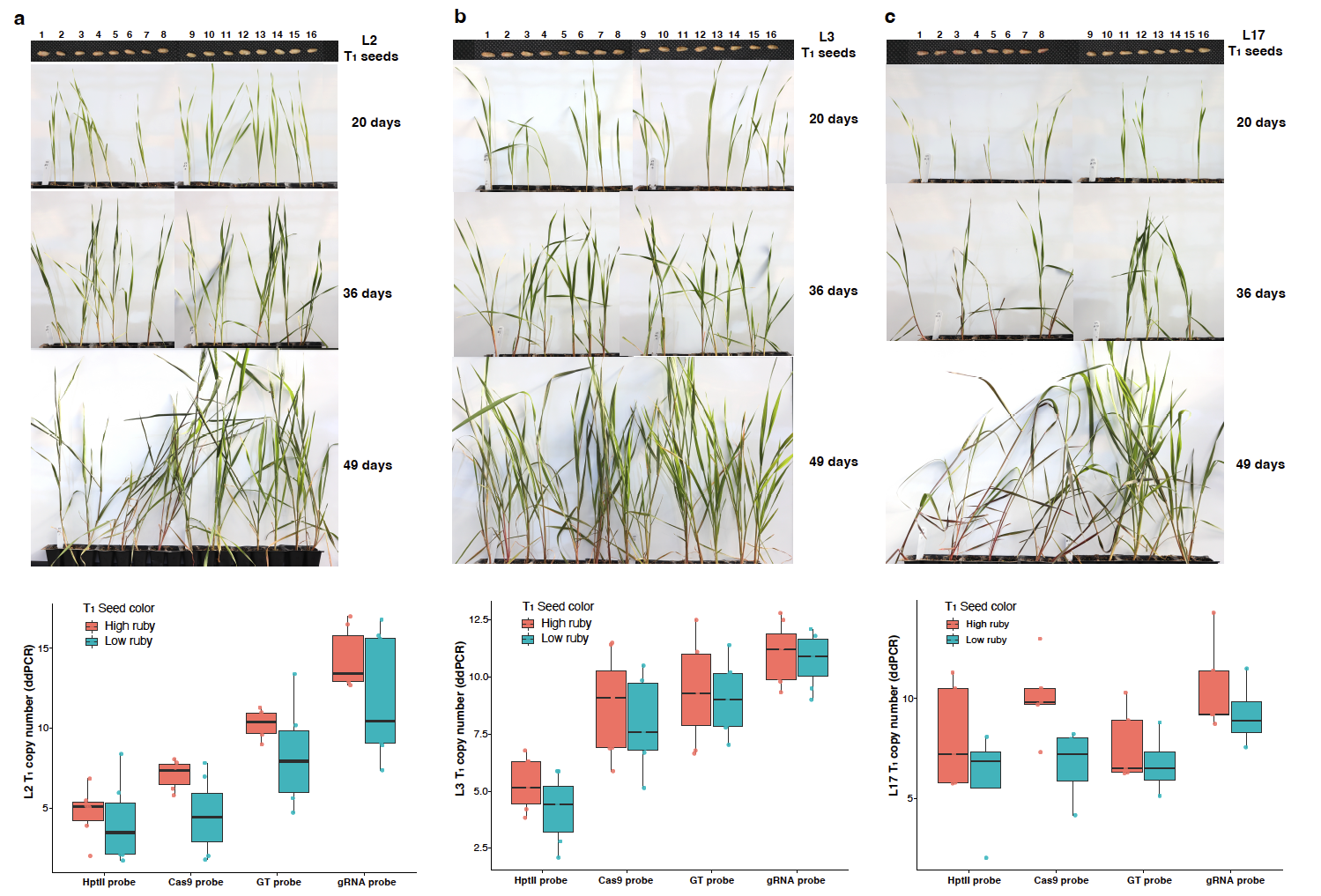

**Figure S3: RUBY-dependent segregation and copy-number variation in two representative T_1_ Fielder wheat lines.** (a-c) Three independent T_1_ lines (L2, L3, and L22) are shown, each derived from a single T_0_ Fielder line. For each line, 16 T_1_ seeds are displayed at the top, grouped by seed-coat ruby intensity into high-ruby (1-8) and low-ruby (9-16) categories. Corresponding seedlings were grown and imaged at 20, 36, and 49 days after sowing, showing betalain pigmentation accumulation in older tissues, particularly in the stems and base of the leaf. The boxplots at the bottom summarize ddPCR-based T_1_ copy-number distributions for *hptII*, *Cas9*, *GT*, and gRNA probes, comparing plants grown from high-ruby (red) and low-ruby (teal) seed pools for each line.
